# Network-Level Associations in Nonlinear Brain Dynamics Predict Transcendent Thinking in a Diverse Adolescent Sample

**DOI:** 10.64898/2026.04.05.716550

**Authors:** Amir Hossein Ghaderi, Xiao-Fei Yang, Mary Helen Immordino-Yang

**Affiliations:** Center for Affective Neuroscience, Development, Learning and Education (CANDLE), Rossier School of Education, Brain and Creativity Institute, University of Southern California, Los Angeles, CA, USA

**Keywords:** Multiscale Entropy, transcendent thinking, functional brain network, nonlinear dynamics, fMRI

## Abstract

Transcendent thinking (TT) is an enduring affective and cognitive process characterized by abstract meaning-making, moral reflection, self-referential integration, and strong emotional engagement. Despite growing interest in its developmental and affective significance, the intrinsic neural dynamics that predict individual differences in disposition to TT remain poorly understood. Most prior work has relied on linear functional connectivity measures, which may be insufficient to capture the nonlinear and multiscale nature of brain dynamics underlying higher-order affective dispositions like TT.

Here, we introduce a nonlinear functional brain network (FBN) framework based on multiscale entropy (MSE) to investigate whether intrinsic resting-state nonlinear brain dynamics predict disposition to TT in adolescents. Functional connectivity was defined as inter-regional similarity in MSE profiles derived from resting-state fMRI, yielding weighted networks that capture scale-dependent dynamical correspondence rather than linear synchrony. Graph-theoretical, spectral, and information-theoretic measures were computed and evaluated against signal-level and network-level null models. Predictive performance was assessed using machine-learning models and compared with conventional time series–based FBNs. Global intelligence (IQ) was examined as a control cognitive variable.

MSE-based network features, particularly spectral energy and Shannon entropy, showed significant associations with TT and enabled reliable prediction of individual differences, whereas time series–based network measures failed to predict TT. No network measures reliably predicted IQ. Overall, these results indicate that intrinsic nonlinear brain dynamics carry predictive information about affective dispositions, rather than domain-specific or network-localized cognitive abilities such as IQ. This work demonstrates that nonlinear, multiscale network representations of resting-state brain activity provide a principled and predictive framework for modeling individual differences in enduring affective dispositions.

## I. Introduction

ENDURING affective-cognitive dispositions are supposed to be associated with large-scale brain network dynamics rather than localized activity within isolated regions [1]. Contemporary affective neuroscience emphasizes that emotional experience and affective states arise from distributed interactions among cortical and subcortical systems, involving the continuous integration of self-referential, sensory, and regulatory processes [1], [2]. From this perspective, enduring affective-cognitive dispositions are not limited to transient emotional responses, but reflect a broader class of integrative brain states that support meaning-making, valuation, and subjective experience. Within this broader affective framework, transcendent thinking (TT) can be conceptualized as a higherorder affective-cognitive capacity that extends basic emotional processing toward abstract meaning, moral reasoning, and a sense of purpose across lived experiences [3]–[6].

Developmental research has increasingly highlighted adolescence as a critical period for the emergence and consolidation of TT. Youths’ disposition to TT has been linked to the maturation and coordination of large-scale functional brain networks [6]–[8]. In particular, recent longitudinal evidence suggests that disposition to TT is associated with increasing interaction between internally oriented systems such as the default mode network (DMN) and executive control networks including the frontoparietal network (FPN) [7]. These findings indicate that transcendent cognition may reflect an advanced form of affective network integration that shapes neural development toward more cohesive large-scale architectures. Despite these advances, it remains unclear whether intrinsic resting-state neural dynamics can predict individual differences in TT [8].

One potential reason for the limited progress in identifying neural predictors of TT may lie in the predominant reliance on linear functional connectivity measures. Most resting-state fMRI studies construct functional brain networks (FBNs) using correlation- or coherence-based metrics [9]–[11]. While such approaches have proven effective for characterizing synchronous interactions and large-scale network topology [12]–[14], they are fundamentally constrained in their ability to capture nonlinear and nonstationary properties of neural signals. Brain activity is increasingly understood as a complex dynamical system, in which higher-order cognitive functions emerge from nonlinear interactions across multiple temporal scales [15]–[17]. Linear connectivity models are therefore likely insufficient to capture the full richness of neural dynamics underlying integrative cognitive–affective processes such as TT.

During collective neural behavior, the brain operates through continuous interactions between bottom-up sensory processes and top-down control mechanisms [13], [18]. These interactions involve dynamic reweighting and reorganization of information across distributed systems, producing nonstationary and nonlinear activity patterns [19]–[21]. Under such conditions, nonlinear connectivity approaches may better characterize delayed, scale-dependent, and higher-order temporal dependencies between brain regions [22], [23]. This perspective suggests that TT, which relies on integrating abstract meaning, self-referential processing, and emotional evaluation, may be more strongly reflected in nonlinear network properties than in linear measures of synchrony.

Multiscale entropy (MSE) has emerged as a powerful nonlinear metric for quantifying the complexity of neural signals across temporal scales [24], [25]. MSE captures the degree of irregularity in a signal after successive coarsegraining, thereby providing a multivalued representation of signal dynamics that spans fine to coarse temporal resolutions. Prior work has shown that reduced neural signal complexity, as indexed by MSE, is associated with cognitive impairment and neuropsychiatric disorders [26]–[28]. Importantly, MSE computed at the regional level has also been linked to large-scale network organization, suggesting that signal complexity reflects not only local dynamics but also the broader network context in which a region is embedded [29]–[31].

These findings raise the possibility that inter-regional similarity in MSE profiles may provide a principled basis for constructing functional connectivity networks that are inherently nonlinear [32]. Unlike conventional connectivity measures that operate directly on time series, an MSE-based approach quantifies similarity between regions based on their scale-dependent complexity trajectories. Regions exhibiting comparable entropy profiles across temporal scales may thus share similar dynamical regimes, even in the absence of strong linear synchrony [32]. Such an approach offers a complementary representation of functional connectivity that emphasizes cross-scale dynamical similarity rather than instantaneous cofluctuation.

In this study, we use a MSE connectivity framework for constructing functional brain networks from resting-state fMRI [32]. For each brain region, MSE is computed across multiple temporal scales, yielding an entropy trajectory that reflects the region’s complexity across different frequency-related modes. Functional connectivity is then defined as the similarity between these entropy trajectories across regions. By shifting the focus from time-domain correlations to multiscale dynamical similarity, this framework is well-suited to capture nonlinear, long-timescale, and nonstationary properties of BOLD activity [24], [25].

We subsequently apply graph-theoretical analyses to characterize both topological and dynamical properties of the resulting MSE-based networks. Specifically, we quantify classical measures of segregation and integration, including clustering coefficient and characteristic path length [33], [34]. In addition, we incorporate a spectral graph-theoretic measure, i.e., network energy, which reflect the stability of synchronization and have been shown to be sensitive to cognitive states and clinical alterations [35], [36]. Finally, we compute Shannon entropy to characterize the overall heterogeneity of connectivity patterns [35], [37]. We used null models to normalize the graph-theoretical measures in order to control for the effects of edge weights [38].

We hypothesize that MSE-based network features will capture nonlinear aspects of functional brain organization that are predictive of individual differences in transcendent thinking. Furthermore, we expect that these nonlinear network measures will outperform conventional time-series–based connectivity metrics in explaining and predicting TT. As a control analysis, we also examine associations with global intelligence (IQ), which is hypothesized to rely more strongly on domain-specific mechanisms rather than whole-brain nonlinear dynamics.

## II. Method

### A. Data and Ethical Compliance

The dataset for this study was obtained from a larger ongoing project that included a wide range of assessments beyond the scope of the present analyses, such as psychosocial interviews, physiological recordings, and neuroimaging procedures (for detailed information, see https://osf.io/gqs34). The research protocol received approval from the University of Southern California Institutional Review Board (UP-12-00206) and followed all institutional ethical standards. Written informed consent was obtained from participants and their parents or legal guardians, as appropriate. All participants received monetary compensation for their participation.

### B. Participants

A total of sixty-five right-handed, typically developing adolescents (36 females, 29 males; ages 14–18 years, M = 15.8, SD = 1.1) took part in this study. They were recruited from public high schools in Los Angeles. Eligible participants were enrolled full-time in school, passing all of their classes, fluent in English, and not subject to disciplinary action either inside or outside of school. None reported neurological or psychiatric conditions, learning or developmental disorders, or histories of substance use or abuse. Students were excluded if they had ever used psychotropic medication, experienced physical or emotional abuse or neglect, or had any medical conditions that would make MRI unsafe. To ensure cultural and ethnic diversity, each participant had at least one parent who was born and raised to adulthood outside the United States, with parental origins spanning 13 countries.

### C. Behavioral Measures

#### 1 Psychosocial Interview Assessing TT

Participants completed an in-depth, two-hour private interview designed to assess TT, following procedures described in [7], and adapted from [39]. Each participant was presented with 40 real-life stories about non-famous adolescents from various cultural backgrounds, representing a broad range of life experiences and emotional themes. These narratives were delivered by the experimenter from memory using a standardized script, followed by a one-minute video segment showing authentic footage of the featured adolescent. Stimuli were presented in Microsoft PowerPoint on a 17-inch Lenovo laptop.

After each story, participants were asked, “How does this story make you feel?” The interviewer then looked down and took notes—ostensibly as a precaution in case of video recording failure, but also to minimize social cues and encourage open-ended, spontaneous responses. All sessions were videorecorded to ensure transcription accuracy.

Videotapes were later transcribed and double-checked for reliability. Independent raters, blind to participant identity, coded responses for evidence of TT based on three primary indicators:

1. Systems-level reflections or moral reasoning: statements that involve evaluating social systems or questioning their functioning (e.g., “It’s strange how being undocumented becomes like a label, limiting what people can do in certain places.”).
2. Broad insights or moral lessons: reflections on universal values, moral implications, or long-term social significance (e.g., “Because children are the future, we must inspire those who will shape society.”).
3. Analyses of character or perspective: inferences about the protagonist’s mindset or internal motivations (e.g., “She realizes she’s not alone—others depend on her.”).

Responses that did not qualify as TT were typically concrete or situational, focusing narrowly on the protagonist’s actions or circumstances (e.g., “I’m glad it worked out for them.”), or reflecting emotional empathy without deeper abstraction (e.g., “I just feel sad for her.”). These were considered context-bound rather than reflective or transcendent.

Each statement identified as indicative of TT received a score of 1. The total TT score for each participant was calculated as the sum of these individual instances across all 40 story trials.

#### 2 IQ assessments

Participants completed two subtests from the Wechsler Abbreviated Scale of Intelligence–Second Edition (WASI-II) (as reported in Gotlieb et al., 2024). These included the verbal and Matrix Reasoning tasks, both administered individually in a quiet, private room by a trained examiner. Raw scores from the two subtests were first converted to age-normed standard scores and then combined to yield each participant’s composite IQ score. One participant completed only the Vocabulary subtest due to time constraints; for this case, an estimated total IQ score was computed based on their verbal score following standard imputation methods. The complete set of psychological and demographic data collected from all participants is publicly available at https://osf.io/9jxaz/.

### D. MRI Data Acquisition

Consistent with the protocol described in [7], participants completed a seven-minute resting-state BOLD fMRI scan during both the first and second sessions, following the psychosocial assessments. They were instructed to stay awake, minimize movement, and allow their thoughts to wander freely. Throughout the scan, a neutral image of a natural landscape—free of any human or animal figures—was presented continuously. To reduce circadian variability, scanning sessions were generally scheduled around midday. Functional images were acquired on a 3T Siemens Trio MRI system equipped with a 12-channel head coil. Functional scans were obtained using a T2-weighted echo-planar imaging (EPI)* sequence (TR = 2000 ms, TE = 25 ms, flip angle = 90°, matrix = 64 × 64, FOV = 192 mm, voxel size = 3 × 3 × 3 mm^3^). Forty-one interleaved axial slices were collected, yielding whole-brain coverage and a total of 210 volumes. High-resolution structural images were obtained using a magnetization-prepared rapid gradient echo (MPRAGE) sequence (TI = 800 ms, TR = 2530 ms, TE = 3.09 ms, flip angle = 10°, voxel size = 1 mm^3^ isotropic, acquisition matrix = 256 × 256 × 176).

### E. MRI Data Preprocessing

Preprocessing of MRI data was conducted using `fMRIPrep` version 20.0.7 [40], which is based on `Nipype` [41] as its workflow engine.

#### 1 Anatomical Data

T1-weighted images were corrected for intensity inhomogeneity using N4BiasFieldCorrection implemented in ANTs 2.2.0 [42]. Skull stripping was performed using the `antsBrainExtraction.sh` script (ANTs), with the OASIS30ANTs template as reference. Segmentation of gray matter (GM), white matter (WM), and cerebrospinal fluid (CSF) was carried out on skull-stripped T1-weighted images using FAST (FSL 5.0.9) [43]. Each participant’s anatomical image was then nonlinearly normalized to the MNI152NLin6Asym template using `antsRegistration` (ANTs 2.2.0), as provided by TemplateFlow.

#### 2 Functional Data

For each resting-state run, preprocessing followed the standard `fMRIPrep` workflow [40]. A reference BOLD image and its skull-stripped version were generated, and susceptibility distortion correction was not applied. The BOLD reference was co-registered to the participant’s T1weighted image using `flirt` (FSL 5.0.9) with a boundary-based registration cost function and nine degrees of freedom. Motion parameters relative to the reference image were estimated using `mcflirt` (FSL 5.0.9) prior to temporal filtering. Slice-timing correction was applied using `3dTshift` (AFNI 20160207). The motion-corrected and slice-time– adjusted BOLD time series were resampled to native space and subsequently normalized to MNI152NLin6Asym space. From the normalized data, confound time series including framewise displacement (FD), DVARS, and global signals from CSF, WM, and whole brain were extracted following the approach described in [44].

#### 3 Denoising and Confound Removal

To minimize motion-related noise, preprocessed BOLD images in MNI space were denoised using ICA-AROMA as implemented in `fMRIPrep` [45], after excluding non-steady-state volumes and applying 6 mm FWHM spatial smoothing. The resulting non-aggressively denoised data were further processed using `XCPEngine`, following the procedures described in Pruim et al. [45]. Temporal filtering was applied using a first-order Butterworth band-pass filter (0.01–0.08 Hz), and mean WM and CSF signals were regressed out. All confound regressors were band-pass filtered within the same frequency range to avoid spectral mismatch.

### F. Functional Connectivity and Network Construction

The brain was parcellated into 264 spherical regions of interest (ROIs) encompassing cortical and subcortical areas, based on a standard atlas [46]. For each region, the mean time series (BOLD signals) was extracted across all voxels and averaged over all voxels. Each parcellated region served as a node in the functional network, and edges were defined as the pairwise time series-correlation or MSE-correlation between regional BOLD signals. Two types of FBN sets were constructed: 1) Time-series–based networks, and 2) MSE-based networks which are described here.

#### 1 Time Series–Based Networks

Time series–based functional brain networks were constructed using linear correlation of regional BOLD signals. Let *x*_*i*_(*t*) denote the mean BOLD signal of region *i* at time point *t*, with *t* = 1, …, *T* .

Functional connectivity between regions *i* and *j* was defined as the Pearson correlation coefficient between their time series:

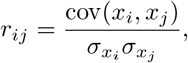

where cov(·) denotes covariance and *σ* indicates the standard deviation of the respective time series. This procedure resulted in a symmetric 264 × 264 weighted adjacency matrix for each participant, with diagonal elements set to zero.

The resulting correlation matrices represented undirected weighted networks, in which nodes corresponded to brain regions and edge weights reflected the strength of linear synchronous coupling between regional BOLD signals. No hard thresholding was applied to the matrices in order to preserve the full distribution of connection weights and to avoid introducing arbitrary sparsity constraints. All subsequent graph-theoretical analyses were performed on these weighted networks.

To assess whether network properties derived from time series–based connectivity reflected information beyond trivial signal characteristics, equivalent adjacency matrices were also constructed from randomly time-shuffled BOLD signals. In these surrogate datasets, the temporal ordering of each regional time series was independently permuted while preserving its amplitude distribution. Equivalent adjacency matrices were generated from the shuffled signals.

#### 2 MSE-Based Networks

MSE-based functional brain networks were constructed by quantifying inter-regional similarity in multiscale entropy (MSE) profiles derived from restingstate BOLD signals. This approach characterizes functional connectivity based on correspondence in nonlinear, scale-dependent signal complexity rather than direct synchronous coupling of time series.

##### a) Multiscale Entropy Computation

For each ROI, the preprocessed BOLD time series *x*(*i*), with *i* = 1, …, *N*, was analyzed using multiscale entropy (MSE). The MSE procedure consists of two steps: temporal coarse-graining and computation of sample entropy (SampEn) at each scale.

At each temporal scale *τ*, a coarse-grained time series *y*^(*τ*)^(*j*) was constructed by averaging non-overlapping segments of length *τ* [47]:

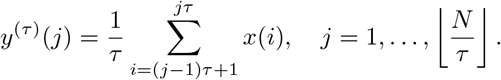

Sample entropy was then computed for each coarse-grained time series. SampEn estimates the negative logarithm of the conditional probability that two sequences of length *m* that are similar within a tolerance *r* remain similar when extended by one additional point [47]:

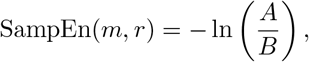

where *B* is the number of template vector pairs of length *m* whose distance is less than *r*, and *A* is the number of matching pairs of length *m* + 1. Following standard practice in fMRI entropy analyses, the embedding dimension was set to *m* = 2. The tolerance parameter was set to *r* = 0.25, which was the lowest value that resulted in less than 5% MSE calculation failure across ROIs. This choice is consistent with previous studies showing that for an embedding dimension of *m* = 2, reliable entropy estimates are obtained for *r* values in the range 0.2–0.4 [32], [48], [49]. These parameters were held constant across regions and participants.

This procedure yielded, for each brain region, a multiscale entropy profile:

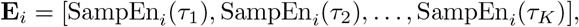

describing signal complexity from fine to coarse temporal scales.

Based on comparisons with randomly time-shuffled surrogate signals, which demonstrated reliable differences between original and shuffled data across scales 1–17, only these scales were retained for subsequent network construction.

##### b Network Construction

Functional connectivity between regions *i* and *j* was defined as the Pearson correlation between their MSE profiles:

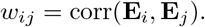

This resulted in a symmetric, weighted adjacency matrix in which nodes corresponded to brain regions and edge weights reflected similarity in nonlinear, multiscale dynamical structure. Regions exhibiting comparable entropy trajectories across temporal scales were therefore considered more strongly connected, indicating correspondence in cross-scale complexity rather than instantaneous signal synchrony.

No thresholding was applied to the adjacency matrices in order to preserve the full distribution of connection weights. Equivalent MSE-based networks were also constructed from randomly time-shuffled BOLD signals using the same entropy and correlation procedures.

This approach yields a weighted, undirected adjacency matrix in which edge weights reflect the similarity of non-linear, multiscale temporal complexity patterns between brain regions, rather than direct synchronous coupling of raw time series. Regions exhibiting similar entropy trajectories across scales were therefore considered more strongly connected, capturing correspondence in cross-scale dynamical organization.

Corresponding MSE-networks were also constructed from randomly time-shuffled BOLD signals by computing MSE profiles and inter-regional correlations using the same procedure. Equivalent adjacency matrices were generated from the shuffled signals.

### G. Graph-Theoretical Network Analysis

Graph-theoretical analyses were applied to all FBNs derived from both time series–based and MSE-based connectivity matrices. All networks were treated as weighted, undirected graphs, with nodes corresponding to brain regions and edge weights reflecting functional connectivity strength. Graph measures were selected to capture complementary aspects of network organization, including modular architecture, global coupling strength, topological segregation and integration, dynamical stability, and informational complexity.

#### 1 Modularity Analysis

The modular organization of functional brain networks was examined using the Newman modularity framework for weighted undirected graphs. In this formulation, modularity *Q* is defined as [50]:

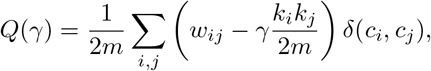

where *w*_*ij*_ denotes the weight of the edge between nodes *i* and *j, k*_*i*_ = ∑ _*j*_ *w*_*ij*_ is the strength of node *i*, 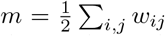 is the total edge weight of the network, *c*_*i*_ indicates the module assignment of node *i, δ*(*c*_*i*_, *c*_*j*_) is the Kronecker delta, and *γ* is the resolution parameter controlling the scale of detected modules.

For a given value of *γ*, modules were identified by maximizing *Q*(*γ*) using the Newman modularity optimization algorithm implemented in BCT. Because the resulting modular structure depends critically on the choice of *γ*, we employed a data-driven procedure to determine an optimal resolution parameter [8], [37], [51].

Specifically, *γ* was incrementally varied starting from the smallest value that yielded more than one module and continuing until the network fragmented into isolated nodes. For each *γ*, modular partitions were obtained and evaluated using a custom optimization procedure [37], [51]. For each identified module, mean within-module connectivity was computed and compared to that obtained from a randomized version of the same adjacency matrix, in which edge weights were independently shuffled while preserving the weight distribution. A modularity ratio (MR) was calculated as the ratio of empirical to randomized within-module connectivity. MR values were averaged across modules to obtain a single summary metric for each *γ* [37], [51].

The value of *γ* that maximized the averaged MR was selected as the optimal modularity resolution. To ensure consistency across participants and network types, this optimization was performed using group-averaged adjacency matrices, and the resulting optimal *γ* was subsequently applied uniformly to all individual networks.

#### 2 Whole-Brain Network Measures

As a baseline descriptor of overall functional coupling, we first computed connection density (CD), defined as the mean weight of all unique edges in the adjacency matrix:

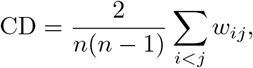

where *n* is the number of nodes and *w*_*ij*_ denotes the weight between nodes *i* and *j*. CD reflects the global level of connectivity independent of network topology.

To characterize network topology, we quantified segregation and integration using the weighted clustering coefficient (CC) and characteristic path length (CPL), respectively. The clustering coefficient measures the extent to which neighboring nodes are interconnected, indexing local segregation. For a weighted undirected network, the clustering coefficient of node *i* was computed following [52]:

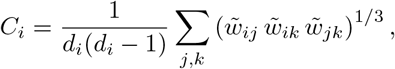

where *d*_*i*_ is the degree of node *i*, 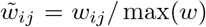 is the weight between nodes *i* and *j* normalized by the maximum edge weight in the network, and the summation is taken over all pairs of neighbors (*j, k*) of node *i*.

The global clustering coefficient was obtained as:

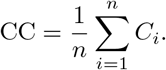

Characteristic path length captures the efficiency of global information integration by measuring the average shortest path between all node pairs. To compute CPL, edge weights were first transformed into distances by taking their inverse, and shortest paths were calculated using the Floyd–Warshall algorithm. Lower CPL values indicate more efficient global integration. To calculate CPL, edge weights were transformed into distances by taking their inverse:

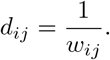

The shortest path length *L*_*ij*_ between nodes *i* and *j* was defined as the minimum sum of distances along all possible paths connecting them. The characteristic path length was then computed as [34]:

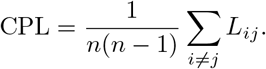

To assess dynamical properties of the networks, we employed spectral graph theory and computed graph energy (*H*), defined as the sum of the absolute eigenvalues of the weighted adjacency matrix [35]:

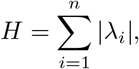

where *λ*_*i*_ denotes the *i*^th^ eigenvalue of the adjacency matrix. Graph energy has been shown to reflect the stability of network-wide synchronization and sensitivity to dynamical state changes in functional brain networks.

In addition, network complexity was quantified using Shannon entropy (*S*), which captures the heterogeneity of connection strengths [35]:

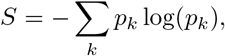

where *p*_*k*_ represents the probability of observing a connection weight within the *k*^th^ bin of the weight distribution. Higher entropy values indicate greater variability and less uniformity in connectivity patterns, whereas lower values reflect more homogeneous network structure.

#### 3 Null-Model Normalization of Graph Measures

To ensure that graph-theoretical measures reflected nontrivial network organization rather than trivial properties such as overall connectivity strength or weight distribution, network-level null models were employed to normalize selected graph measures [38]. Importantly, these null models operate at the level of network topology and are conceptually distinct from the signal-level surrogate analyses based on time-shuffling of BOLD time series.

For each adjacency matrix, null networks were generated by randomly shuffling the connectivity weights while preserving matrix symmetry, network size, and the empirical distribution of edge weights. This procedure disrupts the specific arrangement of connections among nodes while retaining global statistical properties of the network [8], [38]. The null-model randomization was applied independently to all sets of adjacency matrices, including: (i) time series–based networks, (ii) networks derived from randomly time-shuffled time series, (iii) MSE-based networks, and (iv) MSE-based networks constructed from shuffled time-series profiles.

For each empirical network, an ensemble of randomized networks was generated, and the clustering coefficient (CC), characteristic path length (CPL), and graph energy (H) were computed for each null realization. The empirical values of these measures were then normalized by dividing them by the mean of their respective null distributions:

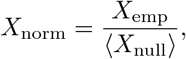

where *X* ∈ {CC, CPL, *H* }.

Shannon entropy (*S*) was not normalized using networklevel null models [8]. Because *S* is defined solely by the distribution of connectivity weights and is invariant to permutations of those weights across the adjacency matrix, shuffling the connectivity weights does not alter its value. Consequently, applying the same null-model normalization to *S* would be uninformative, and the empirical entropy values were retained without normalization.

This normalization strategy ensured that CC, CPL, and H reflected genuine topological and dynamical organization beyond what would be expected from randomized connectivity, while preserving interpretability across different network constructions and surrogate conditions.

### H. Statistical Analyses and Machine Learning

Non-parametric permutation *t*-tests [53] were used to compare the original and randomly shuffled signals. The same procedure was applied to compare MSE values derived from the original signals and those from the randomly shuffled signals across scales 1 to 30. Non-parametric Spearman correlation analyses were conducted to examine associations between normalized network measures and cognitive variables, specifically TT and global IQ. Multiple comparisons were corrected using the false discovery rate (FDR) procedure.

#### 1 Prediction Using Artificial Neural Networks

To investigate whether TT and global IQ could be predicted from network-derived features, we employed an artificial neural network (ANN) with a multilayer perceptron (MLP) architecture with 10 hidden layer nodes. Separate models were trained for each cognitive variable. Two distinct sets of input features were used:

- **Time series-based network features:** Five graph measures (CD, CC, CPL, H, S) computed from adjacency matrices derived from regional BOLD time-series correlations.
- **MSE-based network features:** The same five graph measures (CD, CC, CPL, H, S) computed from adjacency matrices derived from MSE correlations across regions.

The MLP models were trained using Bayesian backpropagation, which estimates posterior distributions over network weights rather than point estimates, enhancing robustness to overfitting and providing uncertainty estimates for the predictions [54]. Model performance was evaluated using 5-fold cross-validation. Prediction accuracy was quantified with the Pearson correlation coefficient (*R*) between observed and predicted scores, as well as the root mean squared error (RMSE). Performance was interpreted as moderate or high when the Pearson correlation coefficient, *R*, exceeded conventional accuracy benchmarks. According to widely used conventions in the behavioral and social sciences, values of *R* around 0.10, 0.30, and 0.50 are typically interpreted as corresponding to small, medium, and large accuracy, respectively [55].

## III. Results

### A. Comparison of Original and Randomly Time-Shuffled Signals

We first compared the original BOLD signals with randomly time-shuffled signals across all 208 time points and cortical regions. Permutation *t*-tests revealed that BOLD signal amplitudes significantly differed between the original and shuffled signals across most time points. Only 28 out of 208 time points did not show a statistically significant difference (Fig. 1(a)). For the MSE values obtained from the original and shuffled signals, significant differences were observed across scales 1 to 17, whereas no significant difference was found for scales 18 to 30 (Fig. 2(a)).

**Fig. 1.**
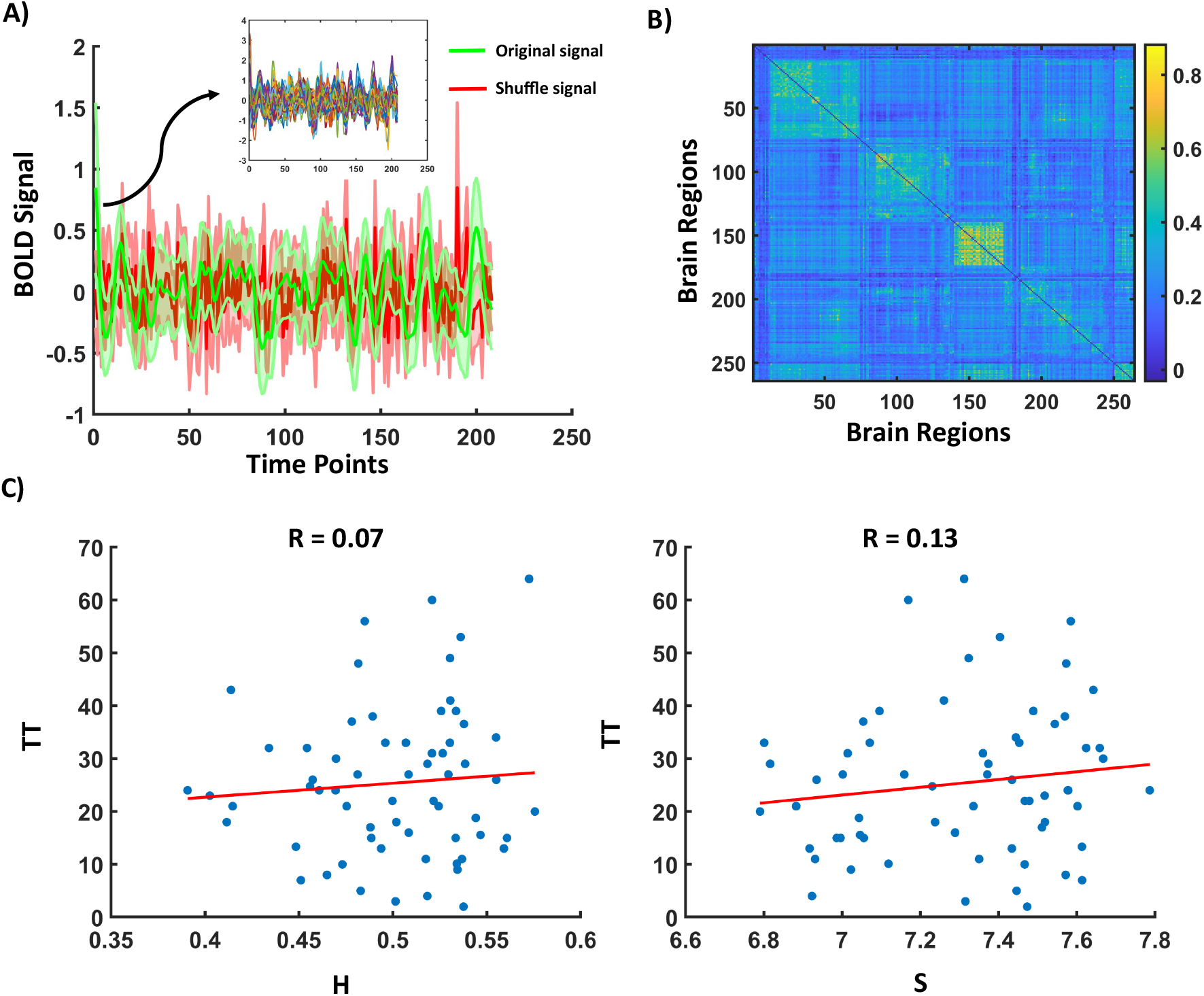
Comparison between original and randomly time-shuffled BOLD signals and their corresponding time series–based functional brain networks (FBNs). (A) Temporal profiles of BOLD signals across time points for original and shuffled data, showing significant differences in most time points. (B) Adjacency matrix of the time series–based functional brain network constructed using Pearson correlation. (C) Associations between time series–based network measures (graph energy H and Shannon entropy S) and transcendent thinking (TT), showing no significant relationships.

**Fig. 2.**
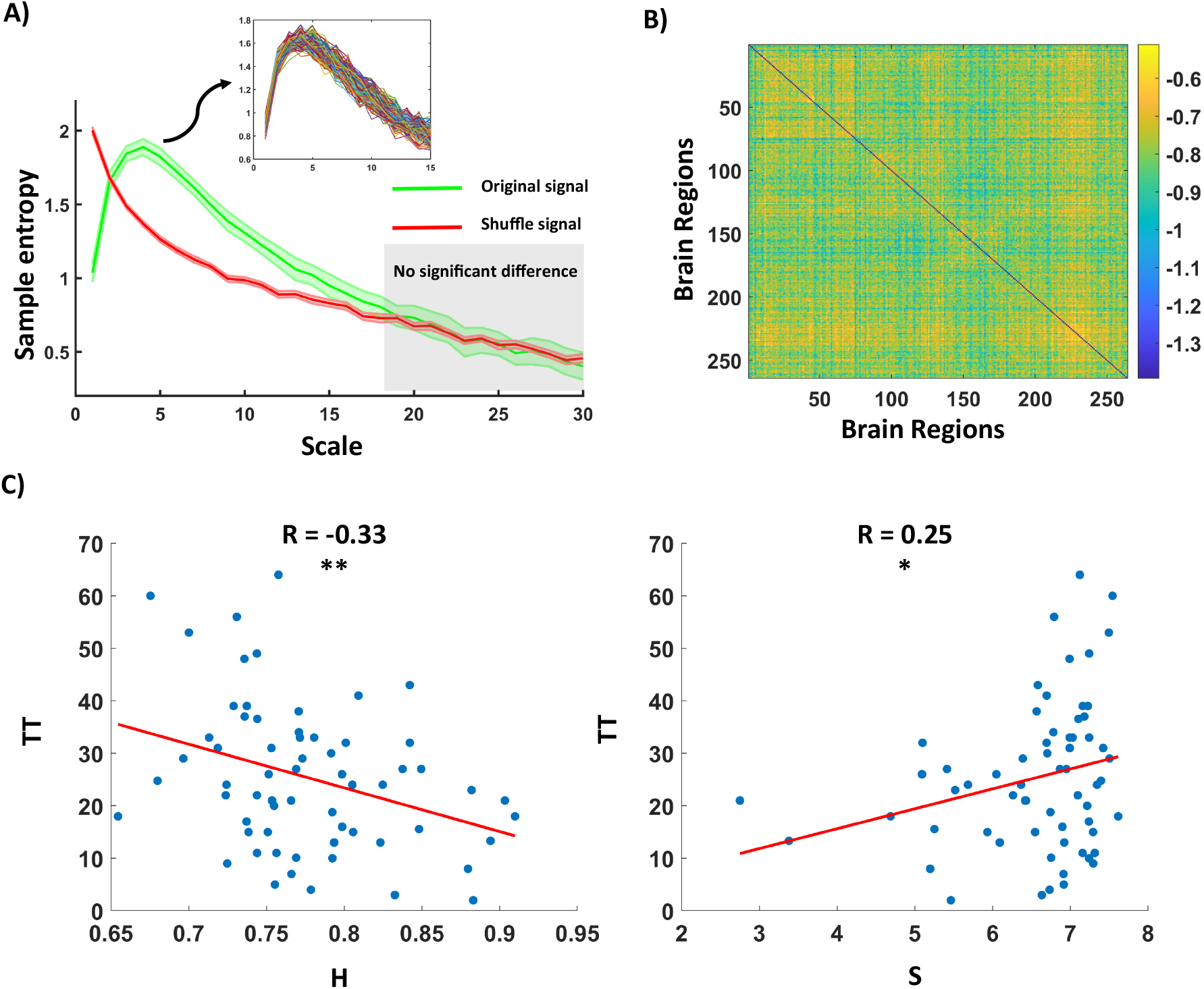
Multiscale entropy (MSE) analysis and MSE-based functional brain networks. (A) Sample entropy as a function of scale for original and randomly shuffled signals. Significant differences are observed for scales 1–17, while no differences appear at higher scales. (B) Adjacency matrix of the MSE-based functional brain network, where connectivity is defined as similarity between regional entropy profiles. (C) Associations between MSE-based network measures and transcendent thinking (TT), showing significant correlations for graph energy (H) and Shannon entropy (S).

### B. Multiscale Entropy Analyses

MSE analyses were performed across all cortical regions for scales 1–17 (fine to mid-range). The scale-dependent trajectory of sample entropy in the original BOLD signals exhibited an initial increase from scales 1 to 5, followed by a decline beyond scale 5. In contrast, the randomly shuffled signals showed a monotonic decrease in sample entropy, with a particularly steep reduction between scales 1 and 5 (Fig. 2(a)). These patterns indicate that the temporal structure of genuine BOLD signals carries meaningful complexity, especially at finer temporal scales.

### C. Network Measures

#### 1 Modularity Analysis

Adjacency matrices for time series- and MSE-based networks are shown in the (Fig. 1(b)) and (Fig. 2(b)), respectively. Modularity analysis revealed that both time series- and MSE-based networks exhibited modular organization. The optimal modular state contained 8 modules for time series-based networks (average *MR* = 1.16) and 17 modules for MSE-based networks (average *MR* = 1.10).

#### 2 whole FBN Measures and Correlations with Cognitive Variables

Five topological, spectral, and entropy-based network measures (i.e., CD, CC, CPL, H, S) were computed for all adjacency matrices. Measures from the original networks were normalized by their corresponding values from the shuffled networks to evaluate deviations from null models. Spearman correlation analyses were then performed between normalized network measures and TT and global IQ, separately. After FDR correction, significant correlations were observed between TT and two MSE-based network measures of H (*R* = − 0.33, *p* = 0.008) and S (*R* = 0.252, *p* = 0.042). In contrast, none of the time series-based network measures was significantly associated with TT, and global IQ did not show significant correlations with any network measure.

### D. Prediction of Cognitive Variables Using ANN Models

ANN models with MLP architectures (Bayesian backpropagation training algorithm, 10 hidden layer nodes) were used to predict TT and global IQ from five graph-based network features. Separate models were trained for time series- and MSE-based networks. Cross-validation indicated high predictive accuracy for TT when using MSE-based network features (*R* = 0.631, RMSE = 11.587, Fig. 3), whereas prediction using time series-based network features did not reach moderate or high accuracy (*R* = 0.082, RMSE = 0.140. Prediction models for global IQ did not achieve moderate or high accuracy with either feature set.

**Fig. 3.**
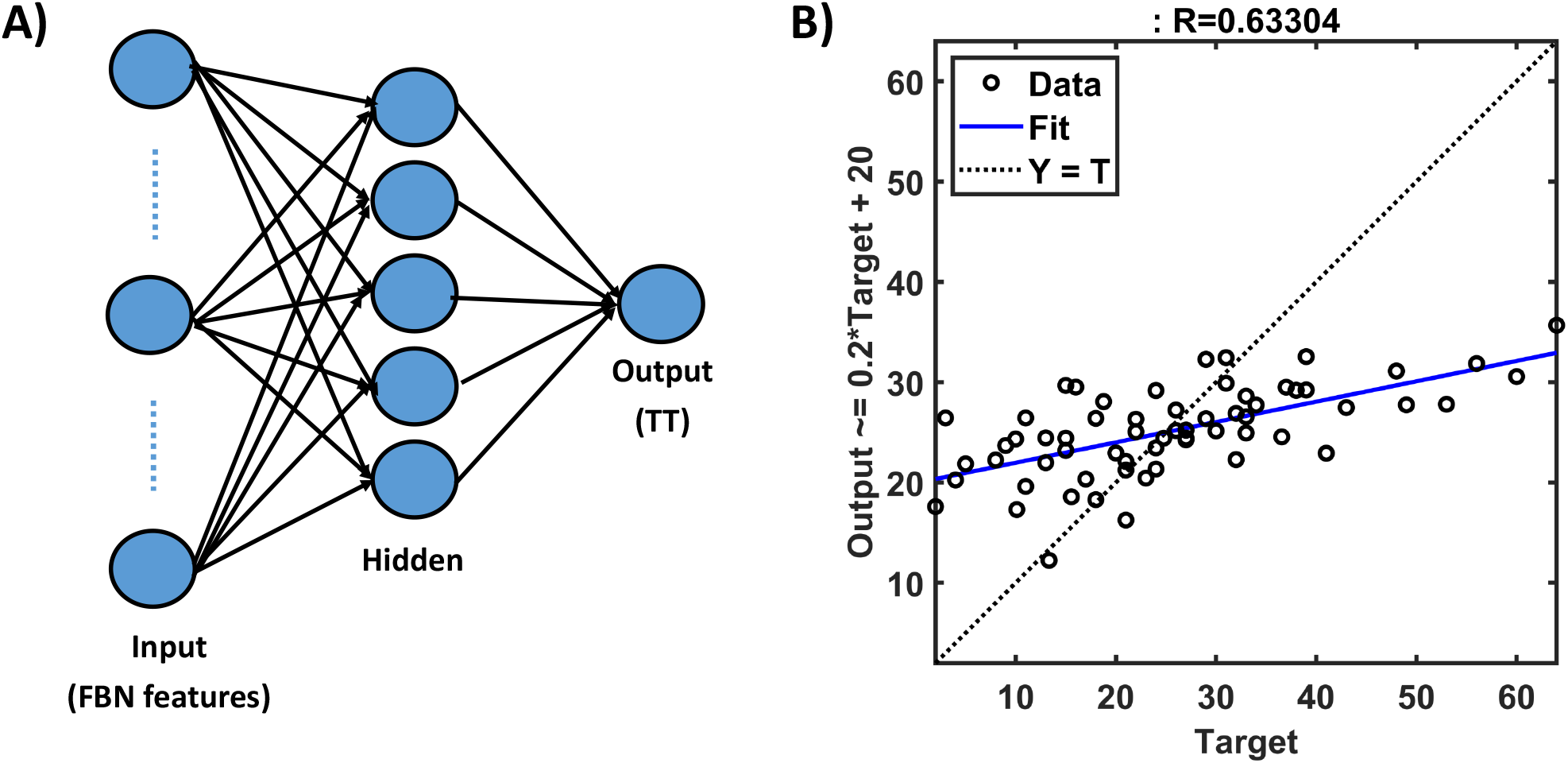
Artificial neural network (ANN) model for predicting transcendent thinking (TT) from functional brain network features. (A) Multilayer perceptron (MLP) architecture with five input features (CD, CC, CPL, H, S), one hidden layer, and TT as the output. (B) Prediction performance for TT using MSE-based network features, showing high predictive accuracy.

## IV. Discussion

The main goal of this study was to compare the efficiency of a nonlinear MSE-network framework with a time-series correlation–network model for predicting TT in adolescents. Before addressing this objective, we conducted a series of analyses to ensure that our methods captured meaningful information. These analyses showed that MSEs across different brain regions exhibited a meaningful multiscale pattern that was significantly different from that observed in surrogate data. The MSE-network produced modular functional networks and showed significant associations with TT, specifically through dynamical measures of graph energy and Shannon entropy. In contrast, no graph-theoretical measures derived from time-series correlation networks were related to TT, and neither connectivity approach showed significant associations with IQ. An ANN-MLP model with Bayesian backpropagation learning further demonstrated that TT could be predicted from MSE-network features with good accuracy, whereas time-series correlation–network features showed poor predictive performance.

### A. Interpretation of Scale-Dependent Entropy Patterns

The scale-dependent trajectory of sample entropy provides clear evidence that resting-state BOLD signals contain meaningful temporal structure. The genuine signals exhibited an increase in entropy from scales 1 to 5, indicating rising complexity at fine temporal scales, followed by a gradual decline at coarser scales. This profile is consistent with nonlinear, multiscale interactions in real neural activity, in which shortrange (high frequency) fluctuations embed additional structure that is not present at longer scales (low frequency fluctuations). This pattern aligns closely with prior reports showing that healthy neural systems exhibit increased complexity at fine temporal scales followed by a progressive decline across coarser scales [24], [26], [29]. This rise–fall trajectory has been interpreted as a hallmark of multiscale coordination in large-scale brain dynamics, reflecting the simultaneous contribution of fast local fluctuations and slower integrative processes [24], [26], [29].

In contrast, the shuffled signals, which preserve amplitude distributions but disrupt temporal dependencies, showed a monotonic reduction in entropy beginning at the smallest scale, with an especially steep decline between scales

1 and 5. This monotonic entropy reduction is consistent with earlier findings demonstrating that disruption of temporal dependencies—through phase randomization, shuffling, or surrogate modeling—eliminates the characteristic multiscale structure of neural activity and yields simplified, noise-like trajectories [**?**], [47]. The pronounced entropy loss at fine scales in the surrogate data replicates prior evidence showing that fine-scale MSE is particularly sensitive to structured neural autocorrelations and cross-scale interactions [25], [56]. Together, these results confirm that the early entropy increase observed in the real signals reflects genuine nonlinear temporal organization rather than amplitude-driven statistical artifacts, reinforcing the suitability of MSE for characterizing neural complexity in resting-state fMRI.

This rise–fall trajectory of entropy across scales was observed across brain regions; however, regions also exhibited meaningful deviations from one another. These similarities and differences reflect region-specific nonlinear and nonstationary dynamics that are not captured by evaluating neural signals solely in the time domain. Consequently, cross-regional comparisons of multiscale entropy profiles provide a principled way to quantify interregional similarity in complex signal structure, thereby motivating the use of MSE-profile similarity as a robust connectivity metric. This observation justifies the subsequent construction of MSE-based FBNs.

### B. Modularity Analyses in Time-Series and MSE Networks

Modularity analysis revealed that MSE-networks exhibited a substantially different organizational pattern compared to conventional time-series correlation networks. While the time-series networks produced eight larger modules with a higher modularity ratio, the MSE-based networks yielded 17 considerably smaller and more differentiated modules with a lower modularity ratio. This divergence suggests that nonlinear multiscale dynamical similarity organizes functional brain networks along dimensions that differ from those captured by linear synchronous activity. This finding is consistent with prior work showing that nonlinear connectivity measures often partition brain regions in ways that do not align with canonical resting-state FBNs identified through correlation or coherence [57]–[59].

In this context, the smaller and more numerous modules detected in the MSE-network likely reflect the ability of MSE-based similarity measures to capture subtle divergences in nonlinear and nonstationary dynamics across brain regions. Because MSE characterizes the complexity of the signal across multiple temporal regimes, it is expected to uncover distinctions that remain latent in linear time-series correlation analyzes. The lower modularity ratio observed in MSE-networks indicates that although regions cluster into meaningful nonlinear similarity groups, these clusters are less sharply segregated than modules derived from time-series correlation. This outcome aligns with previous observations that nonlinear dynamical relationships often produce softer and more overlapping boundaries between subnetworks [57], [58]. Such patterns suggest that nonlinear interactions may support nonlinear distributed and lagged communication rather than strictly synchronous and segregated functional communities. These nonlinear modules may correspond to latent dynamical subnetworks that coordinate cross-scale integration, a form of organization that conventional correlation-based approaches cannot reveal. Thus, the increased number of modules and reduced modularity ratio together highlight that MSE-networks expose a finer-grained and more dynamical brain architecture that is conceptually distinct from traditional functional subnetworks.

### C. Links Between Transcendent Thinking and MSE-Based Network Features

The main finding of this study was that TT was significantly associated with, and could be predicted by, MSE-network properties. In contrast, correlation-based connectivity measures, which quantify only linear synchronous fluctuations between regional time series, failed to capture these relationships. This discrepancy suggests that TT, as a higherorder cognitive–affective construct, is more tightly linked to the brain’s intrinsic nonlinear dynamics than to purely linear functional coupling. Such nonlinear dynamics may include delayed neural responses [22], complex interaction patterns among distributed neural populations [23], [57], [60], coordination across different frequency modes in BOLD signals [61], [62], and the dynamical interplay between local and global network states. These processes can be captured by MSE, where fine scales are thought to reflect more segregated circuits and coarser scales are associated with broader network integration [29]–[31].

More specifically, among all network features examined, two dynamical and information-theoretic properties—graph energy and Shannon entropy—were significantly associated with TT and emerged as strong predictors. In contrast, topological features such as clustering coefficient and characteristic path length showed no significant relationship with TT. This pattern is interpretable in light of the fact that energy and entropy do not capture static topological configurations of the network; instead, they reflect the dynamical stability of synchrony and the diversity of interactions across regions. These findings suggest that TT is not primarily tied to static network topology but is instead more closely linked to the brain’s dynamic regimes.

Taken together, these results indicate that adolescents who exhibit greater diversity in the complexity structure of non-linear interactions in the resting-state brain tend to show more abstract, self-transcendent, and morally engaged interpretations of socioemotional narratives. Conversely, when nonlinear functional brain networks exhibit more stable and less diverse dynamical states, TT tends to decrease. These findings highlight the central role of nonlinear dynamics and multiscale complexity correspondence among brain regions in capturing interindividual variability in higher-order cognitive– affective functioning, emphasizing that such dispositions are better reflected in the brain’s dynamical organization than in its static topological properties.

### D. Lack of Associations Between Network Measures and IQ

Our results indicate that none of the FBN features, whether derived from the MSE-based networks or from the time-series networks, showed a significant association with general IQ. Consistently, the predictive accuracy of our models for IQ was poor. This finding is in line with recent negative evidence suggesting that there is no robust or consistent relationship between general intelligence and global properties of FBNs [63].

Based on this observation, it can be argued that intelligence, as a cognitive trait, is more likely associated with localized and task-specific neural mechanisms rather than with whole-brain network organization. Because the measures examined in the present study were computed at the level of the global functional brain network, they may therefore lack sensitivity as predictors of IQ. This pattern stands in contrast to TT, for which global and integrative properties of functional brain organization appear to provide meaningful explanatory power.

## V. Conclusion

This study demonstrates that nonlinear, multiscale entropy (MSE)–based functional brain networks provide a meaningful and predictive representation of intrinsic brain dynamics associated with TT in adolescence. By defining functional connectivity as similarity in multiscale entropy profiles rather than linear time-series correlations, the proposed framework captures nonlinear and scale-dependent neural organization that is not accessible to conventional connectivity approaches. Importantly, predictive performance was driven by dynamical and information-theoretic network properties, particularly spectral measure of graph energy and Shannon entropy, rather than classical topological measures. This finding suggests that TT is more closely related to the diversity and stability of non-linear network dynamics than to static network architecture.

Overall, these results indicate that intrinsic nonlinear brain dynamics carry predictive information about enduring cognitive–affective dispositions, rather than domain-specific or network-localized cognitive abilities such as general intelligence. TT, as an enduring affective–cognitive disposition that reflects stable tendencies toward abstract meaning-making, moral reflection, and self-referential integration, appears to be embedded in the brain’s intrinsic, resting-state dynamical organization. The proposed MSE-network framework therefore provides a principled approach for linking stable restingstate neural dynamics to individual differences in complex affective cognition and offers a promising tool for predictive modeling of trait-like affective and cognitive dispositions in developmental and affective neuroscience.

## IV. Limitations

Several limitations should be acknowledged. First, the sample size was modest, which may constrain the generalizability of the predictive models despite the use of cross-validation, permutation testing, and null-model normalization. Replication in larger and independent datasets will be necessary to establish the robustness of the observed predictive relationships. Second, the analyses were based on resting-state fMRI, which limits inference about task-evoked or context-specific neural processes. Future studies integrating task-based paradigms may help clarify how intrinsic nonlinear dynamics support predictive processing during explicit transcendent or moral reasoning. Third, multiscale entropy estimation from BOLD signals is constrained by temporal resolution and scan duration. Although reliable entropy scales were carefully selected, higher-temporal-resolution modalities such as EEG or MEG may further enhance predictive sensitivity when combined with the present framework. Despite these limitations, the present findings highlight the predictive value of nonlinear, multiscale brain network dynamics and support their relevance for modeling individual differences in affective and cognitive traits.

